# cblaster: a remote search tool for rapid identification and visualisation of homologous gene clusters

**DOI:** 10.1101/2020.11.08.370601

**Authors:** Cameron L.M. Gilchrist, Thomas J. Booth, Bram van Wersch, Liana van Grieken, Marnix H. Medema, Yit-Heng Chooi

## Abstract

Genes involved in coordinated biological pathways, including metabolism, drug resistance and virulence, are often collocalised as gene clusters. Identifying homologous gene clusters aids in the study of their function and evolution, however existing tools are limited to searching local sequence databases. Tools for remotely searching public databases are necessary to keep pace with the rapid growth of online genomic data. Here, we present `cblaster`, a Python based tool to rapidly detect collocated genes in local and remote databases. `cblaster` is easy to use, offering both a command line and a user-friendly graphical user interface (GUI). It generates outputs that enable intuitive visualisations of large datasets, and can be readily incorporated into larger bioinformatic pipelines. `cblaster` is a significant update to the comparative genomics toolbox. `cblaster` source code and documentation is freely available from GitHub under the MIT license (github.com/gamcil/cblaster).

## Introduction

Complex biological processes are coordinated through the action of multiple distinct, yet functionally associated genes, which are often found physically collocated within genomic neighbourhoods as gene clusters. Gene clusters have been extensively studied in microbes for their ability to encode traits such as the biosynthesis of secondary metabolites, virulence, drug resistance and xenobiotic degradation; however, gene clustering is a phenomenon observed across all kingdoms of life (Chevrette et al. 2020; Wang et al. 2019; Nützmann et al. 2018; Foflonker and Blaby-Haas 2021). Given the relationship between genetic collocation and cofunctionality (Lee 2003; Michalak 2008), it follows that we can detect gene clusters by directly searching for conserved instances of collocation across taxa. Indeed, tools such as MultiGeneBlast (Medema et al. 2013) and clusterTools (Lorenzo de los Santos and Challis 2019) have been developed precisely for this purpose. These tools take specific queries (nucleotide/protein sequences, Hidden Markov Model (HMM) profiles) and search them against genomic datasets, identifying any instances of collocated query hits. However, their utility is limited by the requirement to build and maintain local databases. In many cases, this is redundant given that most biological sequence data is deposited in publicly accessible online databases like the National Center for Bioinformatics Information (NCBI). It is also increasingly demanding as the amount of available data has grown rapidly in the past decade. Between January 2010 and September 2020, the total number of RefSeq genomes grew from 10,171 to 104,969 (NCBI RefSeq Growth Statistics page). This has two additional consequences: i) maintaining an up-to-date local database has become significantly more difficult; and ii) the size of search outputs has increased significantly. Therefore there is a need for search tools that can leverage remote databases and visualise search outputs in an intuitive manner. Here, we present `cblaster`, a Python-based tool to rapidly search for collocated protein coding regions remotely by leveraging NCBI APIs, or locally within user-generated databases. `cblaster` is easy to use with both command line and graphical user interfaces (GUI). It generates fully interactive visualisations implemented in JavaScript and HTML that allow the user to intuit patterns from complex datasets, as well as textual outputs that can easily be incorporated into bioinformatic pipelines. `cblaster` is a significant update to the comparative genomic toolbox and is a launch pad for further functional and evolutionary analyses.

### Fully remote searches against NCBI sequence databases

The `cblaster` search workflow is detailed in Figure 1 and can be launched either through the command line interface (Table 1) or the GUI (Figure 2). A search begins with the user providing either protein sequences (FASTA, GenBank or EMBL format), or a collection of valid NCBI protein sequence identifiers (i.e. accessions, GI numbers), that they believe may form a conserved gene cluster. If the latter is provided, sequences corresponding to each identifier are downloaded using the NCBI Entrez API (NCBI Resource Coordinators 2017) prior to starting the search. Input sequences would typically be taken from the output of a cluster discovery pipeline such as antiSMASH (Blin et al. 2019), or a resource such as the Minimum Information about a Biosynthetic Gene cluster database (MIBiG; Kautsar et al. 2019). Sequences are uploaded to the NCBI BLAST API to launch a new search using BLASTp from the NCBI BLAST+ suite (Camacho et al. 2009; NCBI Resource Coordinators 2017). Optionally, an Entrez search query can be provided to pre-filter the search database, for example to specify a taxonomic group of interest, resulting in vastly reduced search times. Every BLAST search is assigned a unique request identifier (RID) which remains active for 36 hours. A `cblaster` search can be resumed at any point given a valid RID and the corresponding query sequences, such that search results can be retrieved at a later time if desired. This also allows `cblaster` to analyse searches launched through the BLAST website, which can be convenient when expecting long search times due to, for example, many query sequences or unfiltered search databases. `cblaster` can also save search sessions, which can be freely loaded back into the program. The session file contains all data generated by `cblaster` during a search, and can be used to re-detect clusters under new parameters or generate new visualisations without having to completely repeat entire searches.

**Table 1:** Overview of modules provided by the `cblaster` command-line interface.

| Module | Description |
| --- | --- |
| **search** | The main **cblaster** search workflow |
| **makedb** | Create local search databases from personal sequence data |
| **gne** | Gene neighbourhood estimation of **cblaster** search results |
| **extract** | Extract sequences from **cblaster** search results |
| **gui** | Launcher for the **cblaster** graphical user interface (GUI) |
| **extract_clusters** | Extract detected gene clusters from **cblaster** sessions |
| **plot_clusters** | Plot detected gene clusters using **clinker** (Gilchrist and Chooi 2021) |

**Figure 1:**
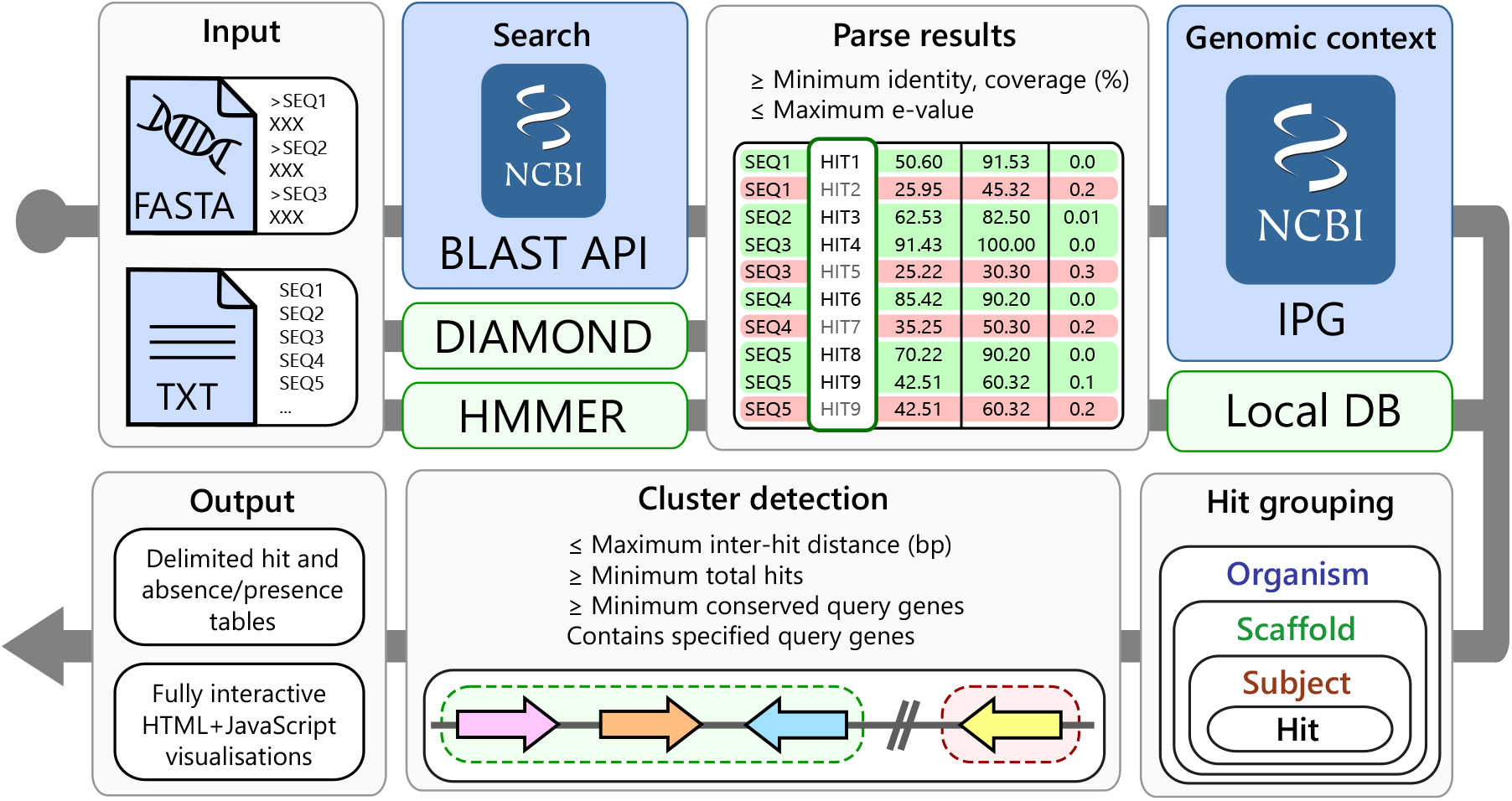
The `cblaster` search workflow. Input sequences are given either as a FASTA file or as a text file containing NCBI sequence accessions. They are then searched against the NCBI’s BLAST API or a local DIAMOND database, in remote (blue background) and local (green background) modes, respectively. BLAST hits are filtered according to user defined quality thresholds. Genomic coordinates for each hit are retrieved from the Identical Protein Groups (IPG) resource. Hits are grouped by their corresponding organism, scaffold and subjects. Finally, hit clusters are detected in each scaffold and results are summarised in output tables and visualisations.

**Figure 2:**
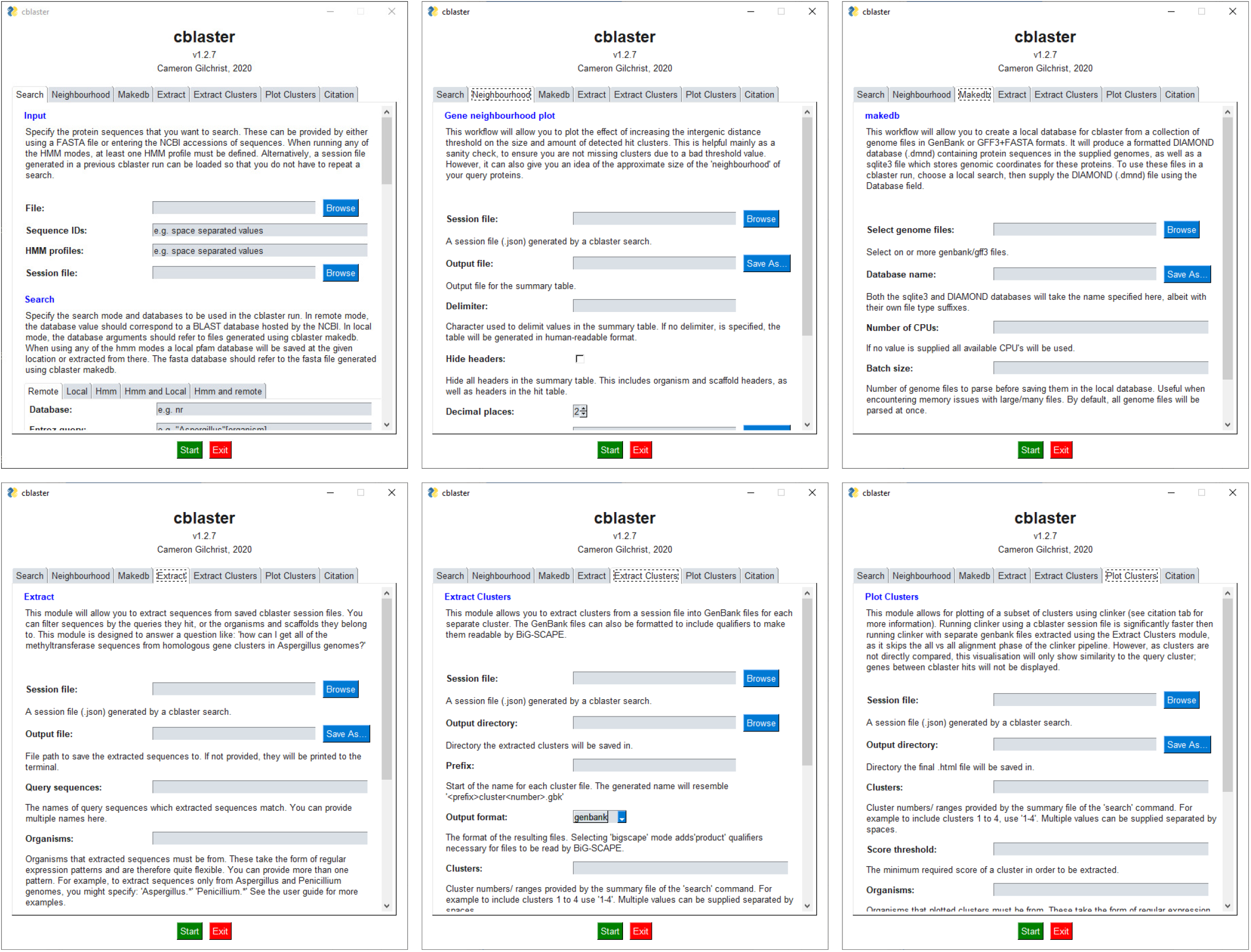
The `cblaster` graphical user interface (GUI). Each panel reprisents a single `cblaster` module: `cblaster search` (top left) for perfoming searches against remote and local databases; `cblaster gne` (top right) for performing genomic neighbourhood estimation; `cblaster makedb` (bottom left) for building local databases from GenBank files and; `cblaster extract` (bottom right) for extracting FASTA files of specific groups of homologues.

The BLAST API is repeatedly polled until the search has completed, at which point search results are downloaded and filtered based on user-defined hit quality thresholds (minimum identity, coverage and maximum e-value). These thresholds are especially useful for narrowing down searches when expecting large result datasets. To retrieve the genomic context of remote BLAST hits, `cblaster` leverages the Identical Protein Groups (IPG) resource via the Entrez API (NCBI Resource Coordinators 2017). The IPG database stores links from proteins to their exact genomic coordinates, meaning BLAST hits can be traced back to their genomic origins. To ensure no potential clusters are missed, for example due to annotation error or fragmented genome assembly, all rows in the IPG table are saved and linked back to their original hits. Hits are grouped by their subject proteins, which are in turn grouped by genomic scaffold and organism. Finally, subjects are grouped into clusters if they contain hits to any required query sequences specified by the user (if any), and satisfy thresholds for maximum intergenic distance, minimum size and minimum hits per unique query sequence. After a gene cluster is finalised, it is assigned a unique numeric ID which is reported in the search output, which can be referenced in other `cblaster` modules.

Clusters are ranked using the scoring formula previously implemented in MultiGeneBlast (Medema et al. 2013). Briefly, cluster similarity is calculated by

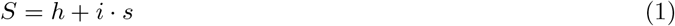

where *h* is the number of query sequences with BLAST hits, *s* is the number of contiguous gene pairs with conserved synteny, and *i* is a weighting factor (default value 0.5) determining the weight of synteny in the similarity score.

### Local searches against custom sequence databases

`cblaster` can also search local sequence databases. The local search workflow mirrors that of remote searches, with two key differences: i) the DIAMOND search tool (Buchfink et al. 2015) is used instead of BLASTp; and ii) a local genome database is used in place of the IPG database to facilitate retrieval of the genomic coordinates of hit proteins.

A local search thus requires two databases to be created by the user: a formatted DIAMOND search database and a local genome database. `cblaster` provides a module, `makedb`, which can rapidly generate both databases from a collection of genome sequences simultaneously in a single command. Briefly, genome files are parsed using the BioPython (Cock et al. 2009) or gffutils (https://github.com/daler/gffutils) libraries. `cblaster` then builds a SQLite3 database containing coordinates of genes within each genome, which can be loaded during local searches. Protein translations are extracted to a FASTA file from which the DIAMOND database is built; sequence headers correspond to unique database indices, allowing `cblaster` to retrieve information about sequences hit in a search.

### Searching for functional composition using domain profiles

Although it is useful to identify clusters through sequence similarity, in some cases this can be too restrictive. Another approach is to search for profile hidden Markov models (HMMs), which correspond to functional domains. In this way, one can search for gene clusters containing hypothetical functionality as opposed to sequence similarity. This is particularly useful when using a ‘retro-biosynthesis’ approach, where biosynthetic gene clusters are identified based on functions informed by chemical structures (Cacho et al. 2015). It can also be useful in cases where mutual sequence similarity between members of a protein family is low. For example, microbial terpene synthases typically show only weak sequence similarity, but can be readily identified through profile HMM searches (Komatsu et al. 2008).

Similar functionality has been implemented previously in ClusterTools (Lorenzo de los Santos and Challis 2019), where users can search local sequence databases for combinations of profile HMMs. `cblaster` provides a search mode, `hmm`, where users can perform searches using profile HMMs from the Pfam database (Mistry et al. 2021) as queries instead of protein sequences. To do this, `cblaster` wraps the the HMMER software package (Mistry et al. 2013). In this workflow, query domain profiles are first extracted from the Pfam HMM profile database using `hmmfetch`, and then searched against a local sequence database (FASTA file of amino acid sequences generated using the `makedb` module) using `hmmsearch`. This requires a local copy of the Pfam database; `cblaster` will automatically download the latest Pfam release if no copy is found. Due to the dependency on the HMMER package, this functionality is currently only supported on Linux and Mac systems. `cblaster` can also perform hybrid searches, where profile HMMs are searched alongside protein sequences.

### Estimation of genomic neighbourhood size

Gene cluster structure can vary greatly between organisms. For example, fungal and bacterial biosynthetic gene clusters tend to be tightly packed, though in plants and some exceptions in fungi, genes encoding biosynthetic pathways are loosely clustered (Kessler et al. 2020; Liu et al. 2020a). In other cases, genes may be split across multiple loci as mini subclusters (Bradshaw et al. 2013), or intertwined with clusters encoding other pathways to form superclusters (Wiemann et al. 2013). During cluster detection, `cblaster` uses a user-defined threshold to determine the maximum distance between any two BLAST hits in a cluster. By default, a cluster is finalised if no new hit is found within 20Kbp of the previous hit. However, this value is arbitrary and may be inappropriate for some datasets, and so it is advisable to test the effect of changing this parameter on each dataset being analysed. `cblaster` provides the `gne` module, which leverages its ability to rapidly reload and recompute search sessions to robustly automate this analysis. When provided with a session file, `gne` iteratively performs cluster detection over a user-determined range of intergenic distance threshold values, then plots the total number of predicted clusters, as well as the mean and median cluster size (bp), at each value (Figure 3b; Additional File 2). These plots typically resemble logarithmic growth, steeply rising at low values but gradually levelling off at higher values. This makes it easy to determine sensible cutoffs that avoid liminal regions where small variability would have a large effect on the output.

**Figure 3:**
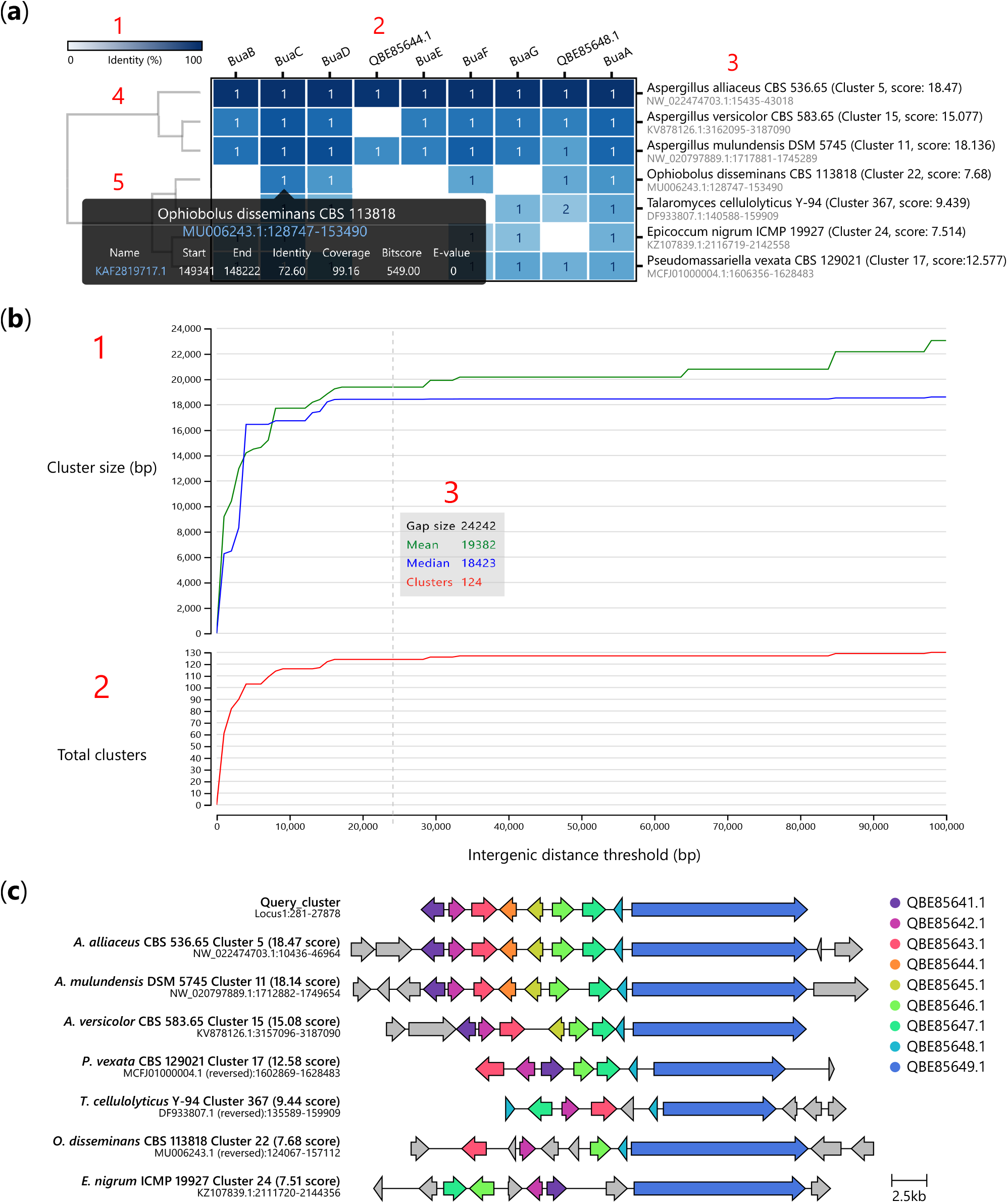
Interactive visualisations generated by `cblaster`. a) Cluster heatmap visualisation of `cblaster` search results: 1) heatmap colour bar indicating 0% (white) to 100% (blue) identity; 2) names of query sequences; 3) names of organism and scaffold locations of hit clusters; 4) dendrogram of hit clusters generated from their identity to query sequences; and 5) cell hover tooltip with detailed hit information including hyperlinks to genomic position on NCBI. b) Gene neighbourhood estimation (GNE) visualisation: 1) Plot of mean and median hit cluster sizes (bp) at different gap sizes; 2) Plot of total clusters at different gap sizes; 3) Hover tooltip showing values of mean and median cluster size (bp) and total clusters at a given gap size. c) Visualisation of gene clusters, inclusive of intermediate genes (grey colour), identified in a) generated using the `clinker` tool via the `plot clusters` module in `cblaster`.

### Comprehensive output and fully interactive visualisations

`cblaster` offers rich summaries and visualisations of search results. `cblaster`’s outputs are available in both human-readable and character-delimited formats, and can be easily incorporated into higher level bioinformatic pipelines. This includes a summary of all detected clusters and a presence/absence table (here termed binary table), which shows the total number of hits in detected clusters per query sequence. By default, clusters are grouped by the organisms and scaffolds they are in; they can optionally be printed in rank order, with the most similar clusters appearing first.

Detected gene clusters can be extracted into GenBank format files directly from `cblaster` search sessions using the `extract clusters` module. Users can filter clusters by their numeric IDs, the organism and scaffold they appear on, or by a minimum cluster similarity score. Optionally, features within each GenBank file can be formatted for interoperability with the biosynthetic gene similarity clustering and prospecting engine (BiG-SCAPE; Navarro-Munõz et al. 2020). In a similar vein, sequences within detected clusters can be extracted using the `extract` module. Sequences can be filtered by the query sequence/s they were hit by during the BLAST search, their organism or scaffold. The `extract` module can produce delimited summary tables as well as FASTA format files of all sequences matching the specified filters.

Search results are visualised as a cluster heatmap (Figure 3a; Additional File 1). Clusters are hierarchically clustered based on best hit identity values using the SciPy library (Virtanen et al. 2020). Cells in the heatmap are shaded based on identity. The text inside each cell indicates if query sequences have multiple hits within a cluster. Figures can be freely panned and zoomed; columns (query sequences) and rows (clusters) can be hidden by clicking on their respective labels. Additionally, mousing over a cell in the cluster heatmap generates a tooltip that displays a summary of hits in the cluster for the corresponding query sequence. The tooltip also links to the exact cluster location on the NCBI’s graphic genome viewer.

Similarly, `cblaster gne` results are visualised as line charts (Figure 3b) which can be panned and zoomed, with a tooltip showing the number of detected clusters, as well as the mean and median cluster size (bp), at different intergenic distance values.

`cblaster` visualisations are implemented as HTML documents that can be opened in the web browser. These documents contain scalar vector graphics (SVG) images generated using the D3 visualisation library (Bostock et al. 2011), which can be exported for modification in vector image manipulation software. Any changes made within the visualisation are reflected in the exported image. Additionally, `cblaster` can produce fully portable HTML documents can be generated, enabling results to be shared between different computers.

Finally, `cblaster` provides a module for analysing subsets of search results using the `clinker` tool (Gilchrist and Chooi 2021), allowing visual comparison of the structure (inclusive of intermediate genes) of specific clusters of interest. This module mirrors the functionality of the `extract clusters` module, whereby clusters can be filtered by ID, organism, scaffold and score.

### Case studies

#### Case Study 1: Analysing evolutionary relationships of chromopyrrolic acid derived natural products

To assess `cblaster`’s ability to visualise evolutionary relationships, we searched for homologues of the *reb* BGC against a local database of characterised BGCs. The *reb* BGC is responsible for the production of the indolocarbazole rebeccamycin in strains of the Actinobacterium *Lechevalieria aerocolonigenes* (Sánchez et al. 2002). Indolocarbazoles are biosynthesised through the successive modification of chromopyrrolic acid, a dimer of of tryptophan. Furthermore, they are closely related to other groups of tryptophan derived natural products, most significantly the indolotryptolines. As such, they represent a good case study to demonstrate the ability of `cblaster` to identify and group related families of BGCs. Using the `cblaster makedb` module, a local database was generated from GenBank sequences downloaded from the October 2019 release of the MIBiG database (Kautsar et al. 2019). The database was queried using protein sequences extracted from the published rebbeccamycin BGC (GenBank accession AJ414559) (Sánchez et al. 2002).

The topology of the resulting dendrogram accurately reflects the previously described groups, indicating a distinction between the indolotryptolines, the rebeccamycins and other indolocarbazoles (Figure 4a, Figure S1). Interestingly, `cblaster` identified two additional groups containing *reb* homologues. Firstly, a group of BGCs encoding structurally unrelated compounds sharing tryptophan halogenases (RebH) and flavin reductases (RebF), including kutzneride (Fujimori et al. 2007), thienodolin (Wang et al. 2016) and ulleungmycin (Son et al. 2017). Although structurally diverse, all members of this group contain a halogenated tryptophan motif in the final structure. Secondly, a group sharing homologues of one or both or the *reb* transport proteins (RebT and RebU), including the glycosides gentamicin (Unwin et al. 2004; Huang et al. 2015) and sisomicin (Hong et al. 2009).

**Figure 4:**
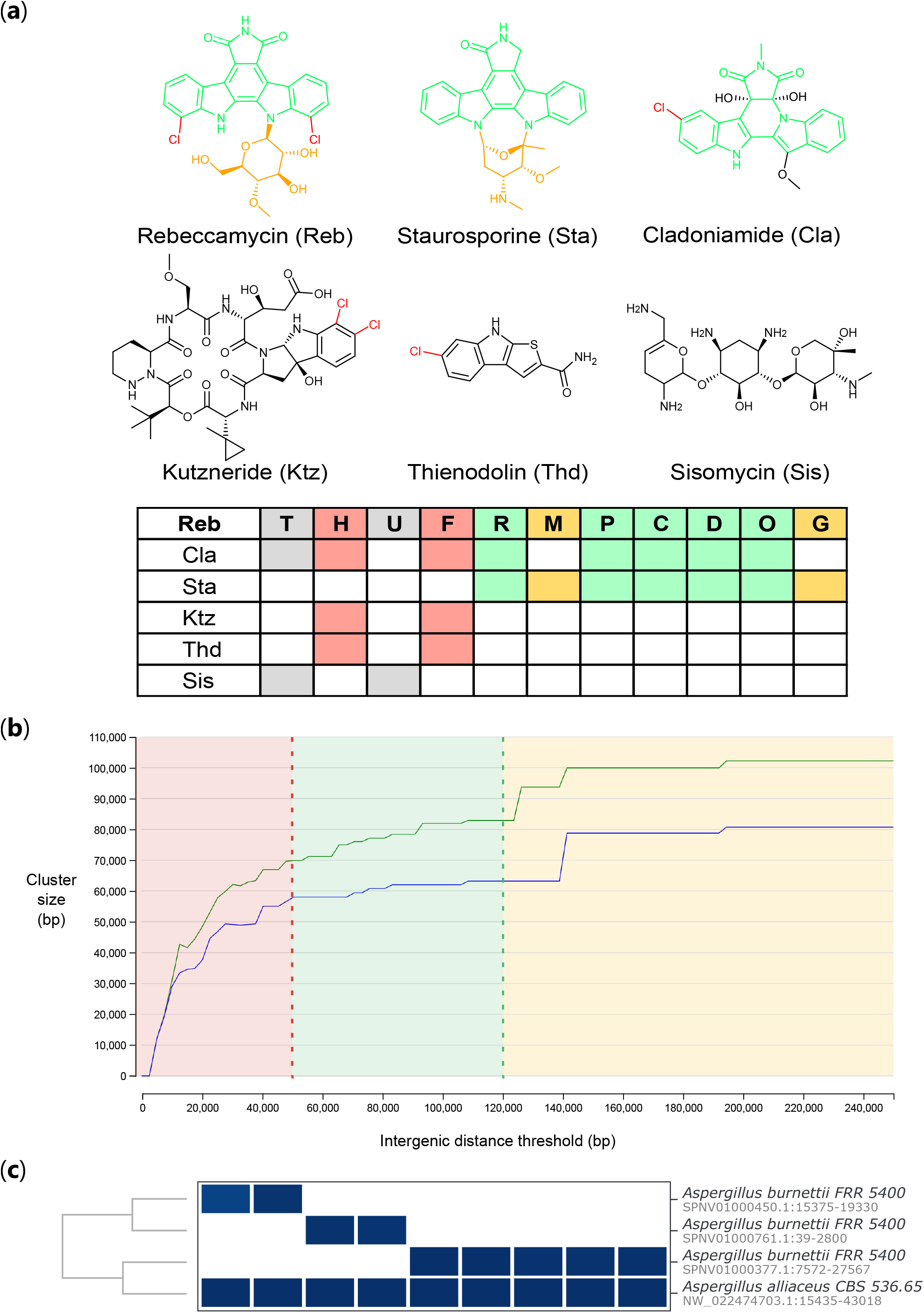
Application of `cblaster` to case studies in bacteria, plants and fungi. a) The evolutionary relationships between rebeccamycin and other natural products as predicted by `cblaster`. Structural features are highlighted according to conserved biosynthetic proteins (halogenases in red, chromopyrrolic acid biosynthesis in green, glycosyltransferase and methyltransferase in yellow). b) Genome neighbourhood estimation of plant triterpene BGCs. ‘Liminal’ region of plot shown in red, ‘stable’ region in green and upper limits in yellow. The accompanying total clusters plot (as in Figure 3b) is ommitted from this figure, but follows the same pattern. c) Using `cblaster` to piece together burnettramic acids BGC in fragmented *A. burnettii* genome.

Although these are only surface level observations, the results demonstrate the ability of `cblaster` to rapidly highlight both apparent and less conspicuous evolutionary relationships.

#### Case Study 2: Genomic neighbourhoods and the biosynthesis of triterpenes in plants

Recent studies have highlighted the importance of genomic neighbourhoods in the evolution of plant development and metabolism (Nützmann et al. 2016; Mihelčić et al. 2019). Brassicacaea for example, have evolved to biosynthesise a range of triterpene products through the process of gene shuffling in plastic regions of the genome (Liu et al. 2020c). During triterpene biosynthesis 2,3-oxidosqualine is cyclised by an oxidosqualine cyclase (OSC). The cyclised product is then decorated through oxidation by cytochrome P450s or acetylation by acetyltransferases. The differing combinations of cyclases and tailoring enzymes drives the remarkable diversity of terpene products.

In most studies, the definition of genomic neighbourhoods is arbitrary, i.e. set to a reasonable, but static value (e.g. selecting five genes either side of the gene of interest). This is analogous to antiSMASH (Blin et al. 2019) prediction where BGC boundaries are set a fixed distance up and downstream of the core genes. We decided to take advantage of `cblaster`’s ability to rapidly recompute outputs to define genomic neighbourhoods in a more robust fashion.

To identify triterpine BGCs, we took the sequences of proteins involved in thalianol biosynthesis in *Arabadopsis thaliana* and searched them against the NCBI database. As described in Field et al. (2011), this included a thalianol synthase (At5g48010), two cytochrome P450s (At5g48000 and At5g47990) and an acyltransferase (At5g47980). We then used the `gne` module to iteratively predict BGCs based on increasing intergenic distance values. From this data, we can readily identify suitable values for the intergenic distance thresholds. Intergenic distances below 50 kb are unsuitable, as they are within the ‘liminal’ region of the curve. Notably, the default threshold of 20 kb lies well within this region, indicating that many BGCs would be missed by `cblaster` if this parameter were not changed. Values above 475 kb leads to a massive spike in the mean cluster size while cluster count remains stable, indicating that homologues are being hit from more distant regions of the genome. Between these two values, there are a number of stable values suitable for analysis, from a conservative threshold of 50 kb to a more liberal threshold of 120 kb (Figure 4b). As such, `gne` allows the user to rapidly assess suitable intergenic distance values that avoid liminal regions where small variability would have a large effect on the output.

#### Case Study 3: Identification of biosynthetic gene clusters in fungi

Previously, we reported the *bua* biosynthetic gene cluster (BGC), responsible for the production of the burnettramic acids in *Aspergillus burnettii* (Li et al. 2019). Due to a fragmented genome assembly, initially we could only identify a short fragment of *bua* containing the genes *buaA* and *buaE*, encoding a hybrid polyketide synthase-nonribosomal peptide synthetase (PKS-NRPS) and proline hydroxylase, respectively. As BGCs are often conserved across multiple species, comparative genomics can be used in this scenario to identify the full set of biosynthetic genes (Cacho et al. 2015). Thus, we employed `cblaster` to search for co-located *buaA/E* sequence homologs across publically available genomes in the NCBI database. This resulted in the identification of complete homologous BGCs in other *Aspergillus* spp., including *Aspergillus alliaceus*, which allowed us to map back to other truncated genomic scaffolds in the *A. burnettii genome* and reconstruct the full *bua* BGC (Figure 4c), which we then experimentally characterised.

In another study, we used `cblaster` to identify BGCs encoding the biosynthesis of natural products with 1-benzazepine scaffolds in *Aspergillus* spp. using genes from the nanangelenin biosynthetic pathway (Li et al. 2020). Searching for collacated homologues of the bimodular NRPS, NanA, and the indoleamine-2,3-dioxygenase, NanC, revealed related BGCs in six *Aspergillus* species. In addition, the different clusters contained different complements of tailoring enzymes, such as methyltransferases and P450s, indicating that these BGCs represented novel analogues of the nanangelenin family.

More recently, we reported *hkm*, the BGC encoding the biosynthesis of the hancockiamides, a family of phenylpropanoid piperazines isolated from *Aspergillus hancockii* (Li et al. 2021). While the BGC itself was identified through manual BLASTp searches of the *A. hancockii* genome, using `cblaster` enabled us to identify homologous gene clusters in several other fungal species.

#### Case Study 4: Identifying drimane sesquiterpenoid BGCs through profile searches

We recently investigated a family of drimane sesquiterpenoids, the nanangenines, isolated from *Aspergillus nanangensis* from section *Jani* (Lacey et al. 2019). Previous work by Shinohara et al. (2016) identified a drimane synthase, AstC, involved in the biosynthesis of the astellolides, a family of drimane sesquiterpenoids. AstC is a novel member of the terpene synthase family and shows similary to haloacid dehydrogenase (HAD)-like hydrolases. The presence of acyl side chains in the nanangenines led us to hypothesize that their biosynthesis would require a drimane synthase, similar to AstC, and either a fatty-acid synthase or PKS. In the study, we performed BLAST searches to identify AstC homologues in the *A. nanangensis* genome, and manually filtered the results for only those found collocated with a PKS or FAS. This led to the identification of the candidate cluster (GenBank accession MT024570.1) that we presented in the paper, containing a PKS (FE257 006541) and AstC homologue (FE257 006542).

`cblaster` provides a search mode, `hmm`, which allows for searches against local sequence databases using functional domain profiles from the Pfam database (Mistry et al. 2021). Our functional hypothesis for the biosynthesis of the nanangenines presented a perfect use case for this search mode. To demonstrate, we attempted to directly identify the same candidate gene cluster in the genome using using the domain profiles for a beta-ketoacyl synthase domain (Pfam accession PF00109) and HAD-like hydrolase (Pfam accession PF13419). This search resulted in just two cluster hits, including our manually identified candidate cluster.

We then performed comparative genomics analysis to further support the validity of our candidate cluster. Though drimane sesquiterpenoids are produced by several Aspergilli, those with acyl side chains are unique to species in section *Usti*. We used `cblaster` to search for homologues of our candidate cluster, both remotely in the NCBI BLAST databases, as well as locally against section *Usti* species with genomes deposited in other sources. As a result, we found homologues spanning up to six genes in *Aspergillus ustus, Aspergillus calidoustus, Aspergillus insuetus* and *Aspergillus pseudodeflectus*, all of which are members of section *Usti* as well as known producers of acyl drimane sesquiterpenoids. Further analyses of these clusters using clinker (Gilchrist and Chooi 2021) gave us confidence that our candidate BGC did indeed encode the biosynthesis of the nanangenines.

Though further studies are necessary to fully characterise this biosynthetic pathway, it shows how `cblaster` has quickly become an indispensable part of this research workflow.

### New approaches and practical applications

The case studies above demonstrate that `cblaster` is a powerful tool for identifying homologous gene clusters from the NCBI or local databases. It is possible to query known gene clusters to expand existing families, or user-specified combinations of genes that are predicted to function together. Currently, the most widely used tools for conducting gene cluster homology searches, MultiGeneBlast (Medema et al. 2013) and clusterTools (Lorenzo de los Santos and Challis 2019), depend on local sequence databases. `cblaster` allows remote searching of NCBI sequence databases, eliminating this requirement and reducing the computational burden (Table 2).

**Table 2:**
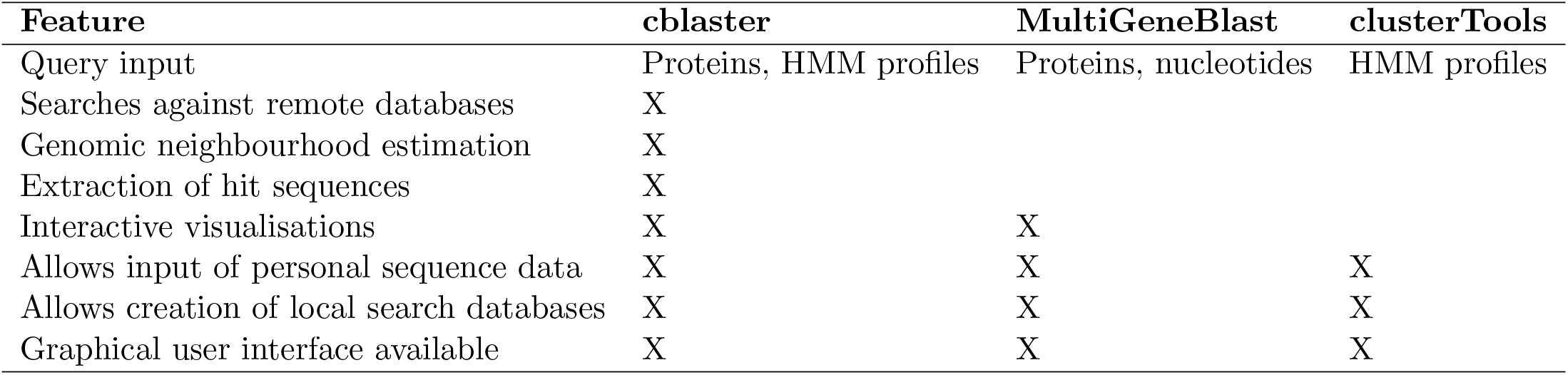
Feature comparison of software tools for gene cluster homology searches.

`cblaster` provides outputs that can be easily adapted for downstream analyses. Delimited formats can be accessed as spreadsheets, making them accessible to all researchers regardless of bioinformatics experience. The binary table can be sorted and filtered post-analysis, which is particularly useful for narrowing large datasets to focus on individual genes or organisms of interest. In this way, data can be easily extracted to generate simple graphs (e.g. generic distribution, ubiquity of different homologues or average cluster size), build distance matrices and dendrograms. Results tables from multiple searches can easily be concatenated to combine results from local and remote searches. `cblaster extract` can be used to quickly obtain specific homologue sequences for downstream phylogenetic and structural analyses.

By using heatmaps as opposed to more complex gene diagrams, information about the diversity and conservation of encoded proteins is accessible at a glance (Figure 3a). Grouping gene clusters by identity highlights blocks of homologous proteins, facilitating the designation of cluster boundaries and identification of core biosynthetic genes and functional sub-clusters. This is a useful proxy for functional and evolutionary relationships, as shown in Case Study 1 above. In this manner, `cblaster` can uncover deep-seated evolutionary relationships that may not be immediately apparent, such as proteins or sub-clusters that have been co-opted between different pathways. Furthermore, `cblaster` can generate comparative gene cluster visualisations using `clinker` (Gilchrist and Chooi 2021). These visualisations show the detected clusters in their genomic context, enabling a fuller view of the overall conservation of cluster architecture across taxa.

`cblaster gne` can estimate the size of genomic neighbourhoods to help evaluate the robustness of cluster prediction (Figure 3b). This enables the user to identify suitable intergenic distance values that limit missing data without inadvertently introducing false positives. In addition, recent work has highlighted the importance of genomic neighbourhoods in the evolution of gene clusters (Liu et al. 2020b). The distribution of genomic neighbourhood sizes, as calculated by `gne`, provides insight into the conservation of gene cluster topology; a broader distribution indicates a more dynamic architecture. This aspect is likely to be of particular interest in the study of plants and other eukaryotes where gene clusters are more variable in gene density, displayed in Case Study 2 above.

In summary, `cblaster` can uncover homologous gene clusters and evolutionary relationships at a pace that far exceeds other available software and has become an indispensable tool in our own research into the biosynthetic gene clusters of fungi and bacteria. It has already aided us in the identification of several metabolic gene clusters from *Aspergillus* species as discussed in Case Studies 3 and 4 above. We foresee `cblaster` being used to uncover novel variants of known pathways, or to prospect for new combinations of biological parts for use in synthetic biology (Wang et al. 2013; Sun et al. 2015; Gilchrist et al. 2018). `cblaster`’s versatility means it can be applied to any project that requires the study of two or more collocated genes and it is accessible to all researchers regardless of their experience with bioinformatics.

### Software implementation and availability

`cblaster` is implemented in Python, with visualisations written in JavaScript and HTML. Source code is freely distributed on GitHub (github.com/gamcil/cblaster) under the MIT license. A comprehensive usage guide as well as API documentation is available online (cblaster.readthedocs.io/en/latest).

## Supporting information

HTML result files for Case Studies 1-4

## Acknowledgements

C.L.M.G. is supported by the Australian Government Research Training Program Ph.D. scholarship. Y-H.C is supported by an Australian Research Council Future Fellowship (FT160100233). This work was funded in part by the Cooperative Research Centres Projects scheme (CRCPFIVE000119).

