## Supplementary material for "cblaster: a remote search tool for rapid identification and visualisation of homologous gene clusters": HTML result files for Case Studies 1-4: CaseStudy1_Output.html

cblaster


cblaster

---

If you found cblaster useful, please cite:

```
						Gilchrist, C.L.M, 2020. cblaster: a Python toolkit for detecting co-located BLAST hits.
					
```

---

Click clusters to hide rows, or queries to hide columns.
The original view can be restored using the reset filters button.

Save SVG
Reset filters


---

Toggle visibility of plot elements:

Hit counts
Multi-hit cell borders
Genomic coordinates


---

Toggle positions of plot elements:

Multi-line cluster information
Query sequence names


---

Adjust size of plot elements:

Cell width:
px

Cell height:
px
