## Supplementary material for "cblaster: a remote search tool for rapid identification and visualisation of homologous gene clusters": HTML result files for Case Studies 1-4: CaseStudy2_Output.html

cblaster


cblaster

This plot shows the effect of increasing the intergenic distance threshold
(--gap argument in cblaster) on both the total amount of predicted
clusters, as well as the mean and median cluster size (bp).

---

If you found cblaster useful, please cite:

```
						Gilchrist, C.L.M, 2020. cblaster: a Python toolkit for detecting co-located BLAST hits.
					
```

---

Zoom with middle mouse, pan by clicking and dragging.

Save SVG
