## Supplementary material for "cblaster: a remote search tool for rapid identification and visualisation of homologous gene clusters": Figure S1

### List of Figures

1 Relationship between the rebeccamycin (*reb*) biosynthetic gene cluster (BGC) and other known BGCs . . . . . . . . . . . . . . . . . . . . . . . . . . . . . . . . . . . . . . . . . . . . . . 2

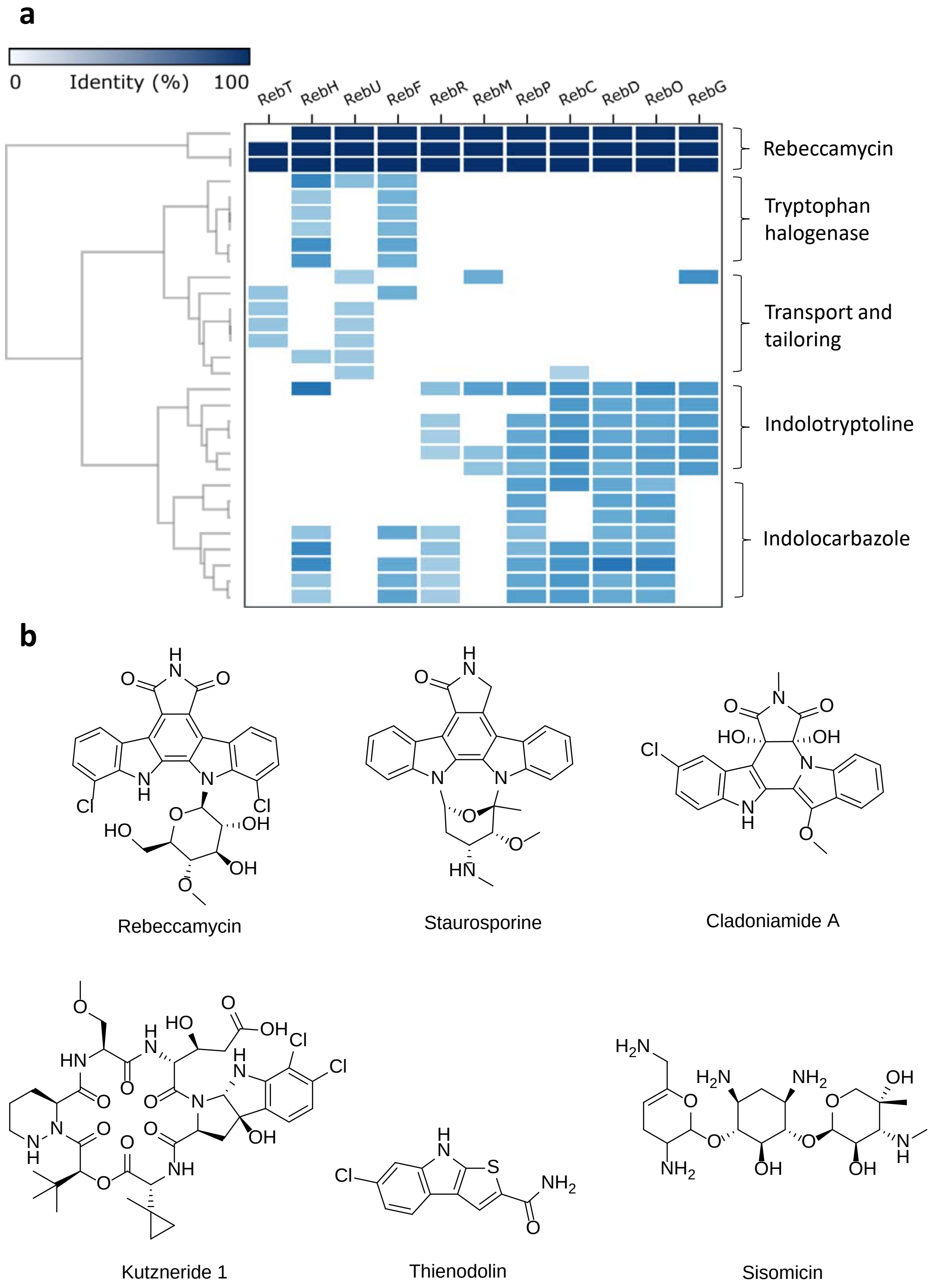


Figure 1: Relationship between the rebeccamycin (*reb*) biosynthetic gene cluster (BGC) and other known BGCs, showing a) the plot as produced by `cblaster` showing the homology of *reb* proteins to those of other BGCs and b) products of rebecamycin and related BGCs as predicted by `cblaster`.
